# Atomic reconstruction of biomolecular structures from AFM images and quantitative validation of experimental data using simulated AFM scanning

**DOI:** 10.1101/2021.06.27.450070

**Authors:** Romain Amyot, Arin Marchesi, Clemens M Franz, Ignacio Casuso, Holger Flechsig

## Abstract

We provide the BioAFMviewer-Toolbox, an extension of our previously developed software platform for simulated AFM scanning of biomolecular structures and dynamics. The focus was on developing a toolbox of methods which employ simulated AFM scanning combined with quantitative analysis to facilitate the interpretation of resolution-limited AFM images. The key advancement is the automatized fitting of biomolecular structures to experimental AFM images, which allows to reconstruct 3D atomistic structures from AFM surface scans. Moreover, several methods for detailed analysis and comparison of surface topographies in simulated and experimental AFM images are provided. We demonstrate the applicability of the developed tools in the interpretation of high-speed AFM observations of proteins. The toolbox is implemented into the versatile interactive interface of the BioAFMviewer, which is a free software package available at www.bioafmviewer.com.

## Introduction

Nowadays nanotechnology allows to observe how single proteins work. Under atomic force microscopy (AFM), e.g., the protein surface can be scanned to image functionally important conformations.^1–5^ The development of high-speed atomic force microscopy (hs-AFM) allowed to even visualize the conformational motions of proteins under near physiological conditions. ^6–10^ While hs-AFM has become a leading technique to study dynamical processes in biomolecules, the analysis and interpretation of the obtained images remains challenging because the experimental spatial resolution limits the visualization of structural details.

On the other side, high-resolution molecular structures of proteins are known^11^ and conformational dynamics can be obtained from multi-scale molecular modelling.^12–14^ The enormous amount of available high-resolution protein data offers the great opportunity to better understand resolution-limited AFM scanning experiments. As an important step in that direction, we have recently developed the BioAFMviewer software platform which provides an interactive interface for simulated AFM scanning of biomolecular structures and conformational dynamics.^15^ The simulated scanning method computationally emulates scanning of the molecular structure to produce pseudo AFM images. It has been employed in several studies to interpret experimental AFM images of proteins, ^8,16,17^ and in molecular simulations to perform flexible fitting of biomolecular structures to experimental images^18^ or to deduce information on the AFM tip shape.^19^

The strengths of the stand-alone BioAFMviewer software are its user-friendly versatile interface and rich functionality. The visualization of molecular structures and simulated scanning to display corresponding AFM graphics proceeds in a synchronized way, even for very large protein structures. Scanning parameters such as tip-shape and spatial resolution, as well as the displayed range of topography heights, can be conveniently adjusted. Furthermore, molecular movies of conformational motions can be processed, which in principle allows simulated AFM experiments of functional dynamics in biomolecules.

With the BioAFMviewer-Toolbox we report here an important software extension. The main purpose was to develop a procedure for automatized fitting of biomolecular structures to AFM images, aiming to predict 3D atomistic structural information from resolution-limited AFM surface scans. Various tools which allow detailed analysis and comparison of surface topographies in simulated and real AFM images, aimed to provide further validation of experimental observations, were also implemented.

## Results

Our report here is focused on the presentation of new BioAFMviewer toolbox developments, and on demonstrating their applications to experimental AFM images. A detailed explanation of the simulated AFM scanning method and on general BioAFMviewer applications is provided in our previous report.^15^

### Fitting biomolecular structures to AFM images

The core improvement of the BioAFMviewer is a toolbox which implements optimal fitting of high-resolution biomolecular stuctures to resolution-limited experimental AFM images. The purpose of this application is to extract information on the three-dimensional molecular structure from just the surface representation available from real AFM images.

Generally, fitting of a molecular structure to an AFM image can be viewed as a complex optimization problem of identifying, from the pool of all possible conformations, a subset for which simulated and experimental AFM image match best. The interactive BioAFMviewer interface, shown in Figure 1, allows the straight-forward design and implementation of a simplified fitting procedure. The synchronized live scanning of the instantaneous orientation of a biomolecular structure displayed in the molecular viewer allows us to directly compare (in principle) *all possible* corresponding simulated AFM graphics to the target experimental AFM image uploaded by the user. The fitting strategy should therefore aim to efficiently sample the space of possible rigid-body orientations of the loaded biomolecule, score the similarity of the corresponding simulated AFM to the user-defined target AFM image, and select candidates with show the best match.

**Figure 1:**
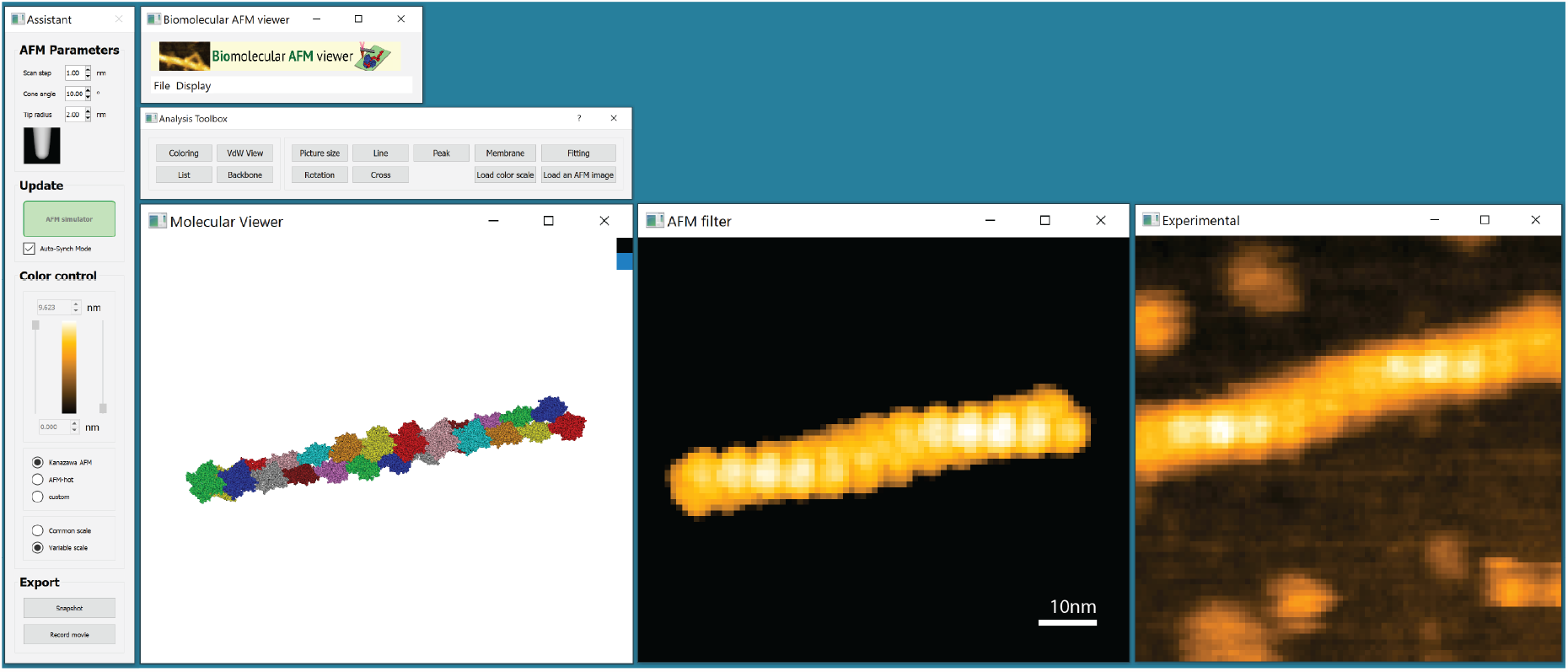
BioAFMviewer interactive interface. Central are the synchronized molecular viewer and AFM windows which provide live simulated scanning of the loaded molecular structure in arbitrary orientations. The *Assistant* window (left) allows to conveniently change scanning parameters such as spatial resolution and tip shape, and to adjust the height range and color scale of the displayed simulated AFM graphics. The *Toolbox* provides the implemented topography analysis tools and the tool for fitting to an uploaded experimental AFM image (right).

We developed a procedure which employs two search strategies to identify molecular structures which best match the target AFM image. A method of *Global Search* was based on extensive unbiased structure sampling, while the *Local Search* methods provides a more sophisticated search algorithm for refined optimization.

### Global search strategy

The *Global Search* method relies on unbiased sampling orientations of the loaded structure in the molecular viewer. The search space was defined by rigid-body rotations around all three spatial axes within the coordinate system of the molecular viewer in discrete steps (see Methods). Similar to the usual workflow of the BioAFMviewer, each sampled molecular structure undergoes simulated AFM scanning to compute an AFM image that can be compared with the experimental target image. A variety of scoring functions is provided to quantitatively evaluate their similarity (see Methods). After completed fitting the user can conveniently access results in the *Fitting Window*. The five molecular structures with the highest scores are presented together with their corresponding simulated AFM images. The reason to provide not only the best matching structure with the highest similarity score as the single optimal fit, is that it may not necessarily represent the visually best comparison to the target AFM image. This is mainly because of the limited spatial resolution of AFM images and the approximate method of simulated scanning, which generally prevents a unique result that can be granted full trust. Particularly for proteins with symmetric domain arrangements ambiguities can be expected. We therefore provide a candidate choice from top fits.

### Local search - Quick Fit function

While the global search function samples molecular structures along a grid of globally available orientations, we also aimed to also provide a more sophisticated fitting method employing a refined search strategy. The Quick Fit function performs a local search of rigid-body orientations confined to the neighborhood of a given initial molecular orientation, and identifies a single molecular structure which best fits the target AFM image. We applied a computationally efficient method which resembles a simplified Metropolis algorithm of optimization (see Methods). The initial molecular orientation can be, e.g., the result of a search by hand, during which the simulated AFM image of the instantaneous structure displayed in the molecular viewer is visually compared to the target experimental image. We have previously demonstrated that manual search can efficiently reveal atomic protein structures from experimental hs-AFM images.^15^ On the other hand, the initial orientation may correspond to a candidate fit obtained from global search. In both situations, the Quick Fit function can be used to obtain a further refinement of structure fitting.

### Application to experimental AFM images

To demonstrate the implemented algorithms of optimal fitting, we applied it to reconstruct the atomistic 3D molecular structure of the cyclic nucleotide-gated ion channel SthK from the observed AFM surface scanning images. Hs-AFM imaging of this channel reconstituted into bilayer membranes revealed the formation of 2D lattices.^20^ We considered an AFM image obtained from scanning the extracellular side of the channel lattice shown in Fig. 2A (see SI for details). For a set of 15 different SthK surface scans within a selected array (marked in Fig. 2A) we individually performed optimal fitting of the atomistic structure of SthK (see Methods). To determine the optimal match of simulated AFM image to the individual experimental target image, a combination of *Global Search* using a coarse searching grid, and refined *Local Search* was always employed (see Methods). In Fig. 2B simulated SthK images after completed optimization are shown. They compare remarkably well to their respective experimental AFM target images, quantified by very high overall comparison scores (see SI). The corresponding atomistic structures are shown in Fig. 2C.

**Figure 2:**
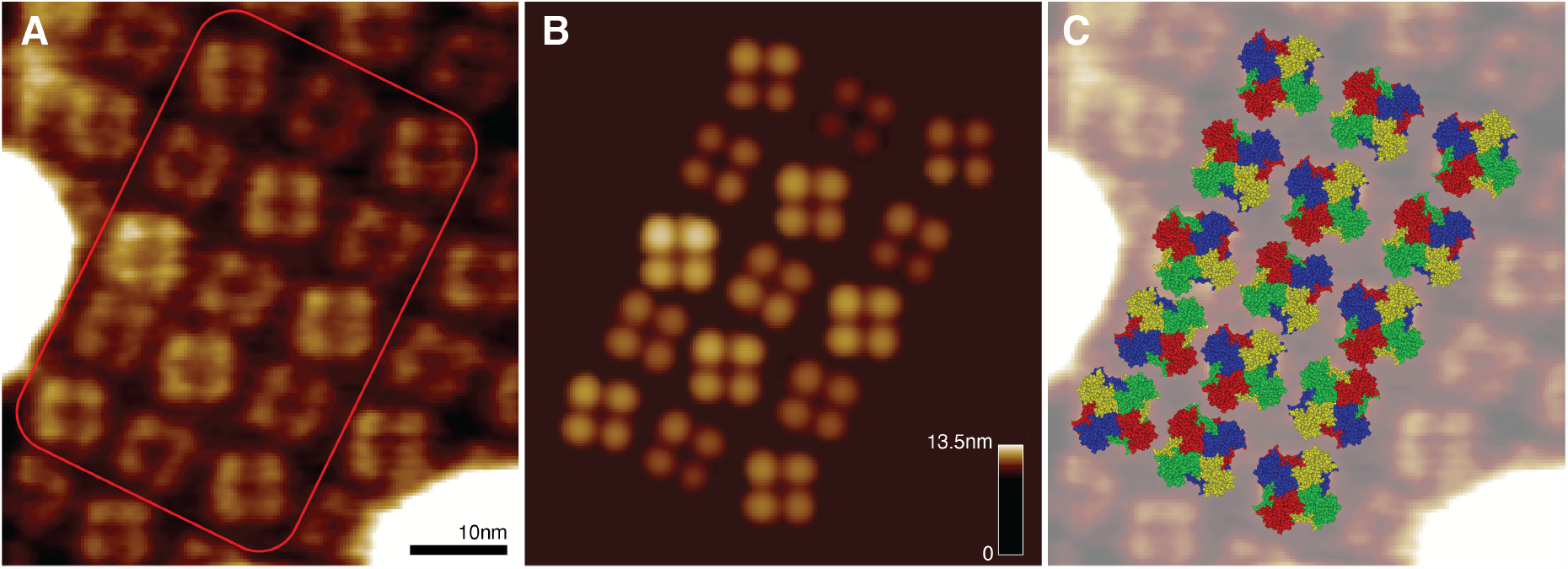
Optimal fitting to AFM images. A: Hs-AFM image of a 2D lattice formation of SthK channels produced from scanning the extracellular side. B: Simulated AFM images obtained from optimized fitting the molecular channel structure individually into 15 selected AFM surface scans (marked in A). C: The array of corresponding atomistic SthK structures superimposed with the experimental AFM image (transparent). Colors represent different protein chains.

This is a strong demonstration of how the BioAFMviewer fitting function can be employed to predict atomistic molecular conformations from AFM surface scans. In Figure 3 further applications are provided, including fitting of the actin-filament all-atom structure to an experimental Hs-AFM image (see SI for details), and those of the ClpB chaperone and F1-ATPase motor which we have previously considered in our first report of the software.^15^

**Figure 3:**
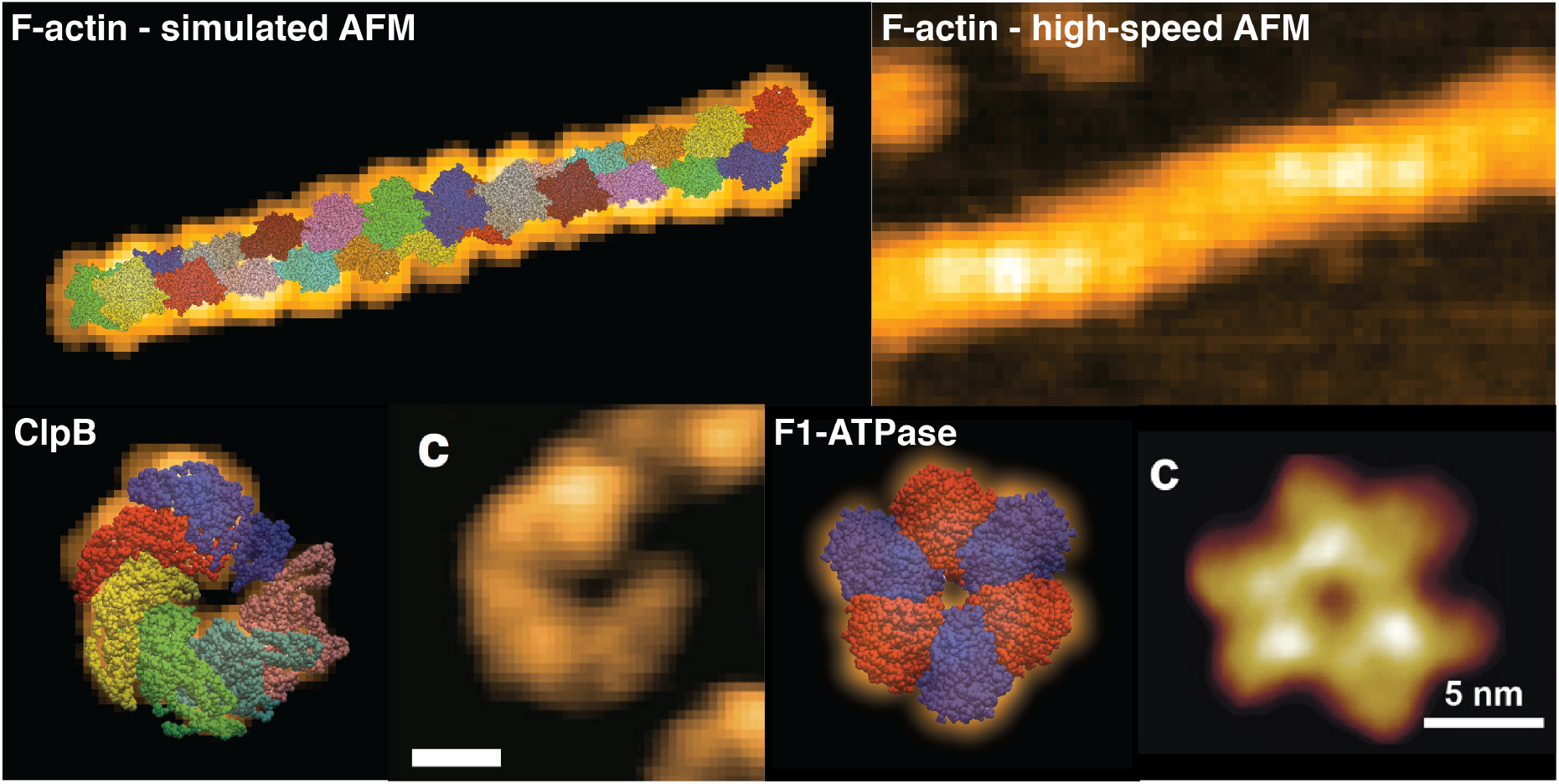
Fitting to AFM images. Top row: Superposition of the actin-filament all-atom structure and the simulated AFM image (left) obtained from fitting to the experimental Hs-AFM image (right). Bottom row: Reconstruction of the atomistic protein structure from fitting to AFM images for the ClpB chaperone (left), and the F1-ATPase (right). The ClpB Hs-AFM image is adopted from^16^ and that of F1-ATPase is from.^10^

### Topography analysis tools

Image comparison methods provides a quantitative assessment of the overall similarity between simulated and experimental AFM images. Beyond that, it is also desirable to have tools available which allow a comparison on a more detailed level, e.g., those analyzing the height topography of the scanned biomolecular surfaces.

We have therefore implemented the *Line tool, Cross tool*, and *Peak tool*, which all enable the user to conveniently select specific image regions for which topography analysis shall be conducted.

A demonstration of topography tools is provided in Figure 4. The *Line Tool* allows the user to draw an arbitrary line in the window canvas of the simulated AFM graphics and that of the uploaded AFM image. The height topography of the scanned molecular surface along the chosen line is then computed and displayed. As a demonstration, this tool was applied to analyze surface topographies of filamentous actin (F-actin) obtained from simulated scanning (Fig. 4A) and from experimental high-speed AFM observations (Fig. 4B, see Appendix). Measuring the consecutive arrangement of filament incorporated actin monomers (line 1) shows a very well agreement of simulated and experimental images, both revealing a spatial separation between neighboring monomers units of about 5.4nm (Fig. 4C). Comparing the line profiles obtained over larger distances (line 2) also shows a very well agreement of the scanned periodic arrangement along the filament structure (Fig. 4D), with the pitch measured as ∼ 35nm and ∼ 32nm. This is compatible within the evident approximations of simulated scanning, and further taking account that in the experiment the actin filament was stabilized by phalloidin molecules which may alter the native structure.

**Figure 4:**
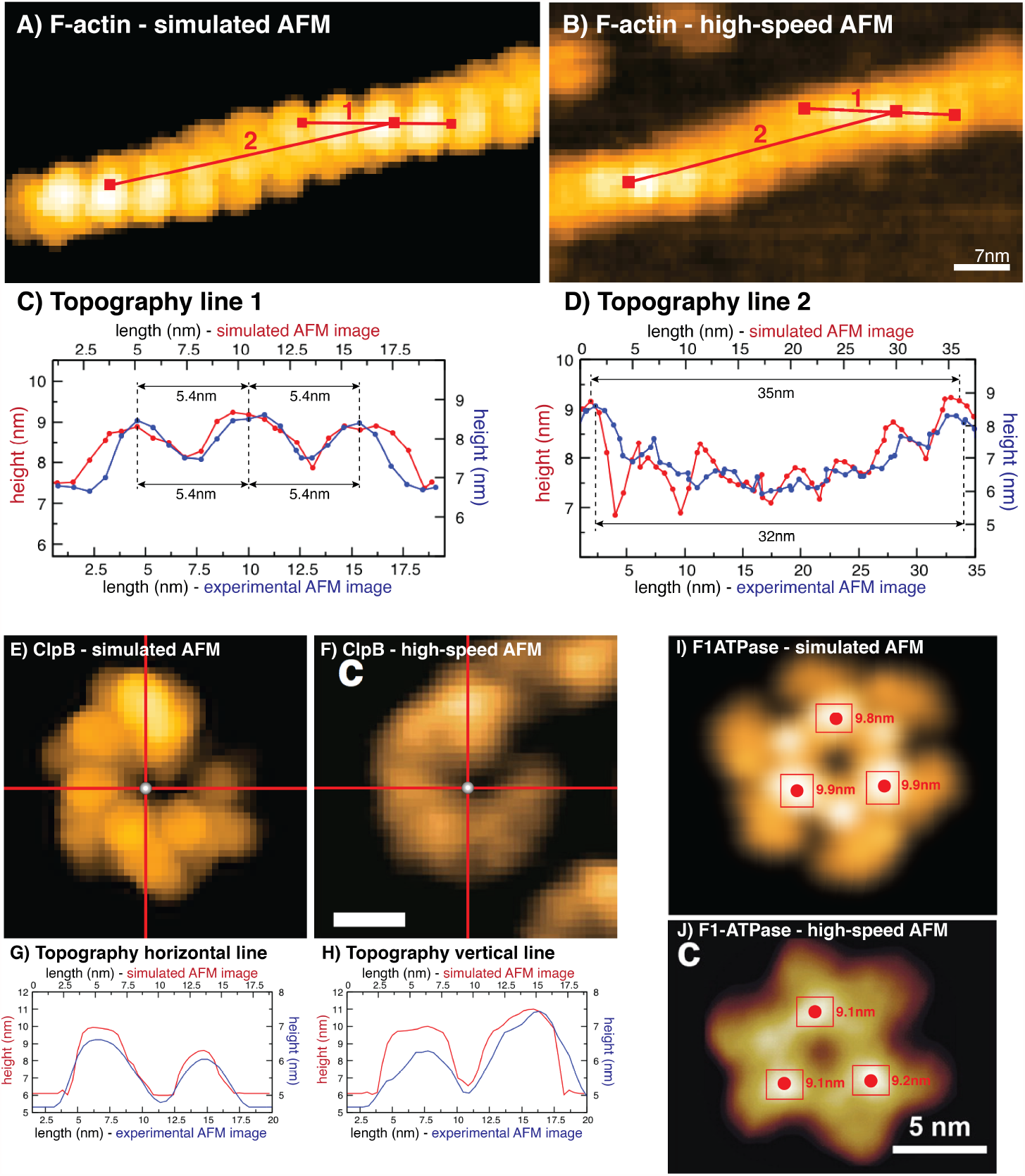
Demonstration of the Topography Tools. A-D: Line Tool function applied to the simulated and experimental AFM image of F-actin (A,B). The two chosen lines are displayed in red. Their corresponding height topographies obtained from simulated and AFM scanning of the molecular surface are compared in panels (C) and (D). E-H: Cross Tool applied to the ClpB protein. Height profiles along vertical and horizontal red lines chosen in simulated (E) and experimental AFM images (F, adopted from^16^) are compared in panels (G) and (H). I,J: Peak Tool applied to F1-ATPase images. In both simulated (I) and AFM graphics (J, adopted from^10^), user-selected regions are marked by red rectangles, inside which red spheres indicate positions with the largest surface height (corresponding values are given).

To demonstrate the *Cross Tool* and *Peak Tool* functions we consider examples from our previous publication^15^ and now provide further quantitative analysis of fitted simulated images to hs-AFM images. The *Cross Tool* allows the user to navigate the origin of a cross formed by horizontal and vertical lines along the canvas of simulated AFM and experimental AFM window, upon which the profiles of scanned surface heights along the corresponding cross sections are computed and displayed. Figure 4 (E,F) presents this tool applied to simulated and hs-AFM image of the ClpB chaperone protein.^16^ The height profiles given in Figure 4 (G,H) show a well agreement of relative surface heights in simulated and AFM images. Finally, the *Peak Tool* allows to specify rectangular shaped regions in the canvas of the simulated AFM and experimental AFM window, inside which the location of the highest protrusion value is detected. Application of the *Peak tool* to simulated and hs-AFM images of the nucleotide-free conformation of the rotorless F1-ATPase protein motor^10^ (Figure 4I,J) show a very well agreement.

Although we have already previously discussed the limitations of simulated scanning,^15^ we find it worth to note how they appear in the performed quantitative analysis of AFM topographies when the comparison to hs-AFM images is undertaken. The main aspect is that simulated scanning, based on non-elastic collisions of the tip with the molecular surface, apparently produces surface heights of systematically larger magnitude compared to experimental AFM scanning which is certainly more complex and perturbative. Apart from that, the agreement of relative heights of the scanned proteins is remarkable.

### Membrane tool

AFM is often applied to study conformational dynamics in proteins which operate inside membranes, such as transporters or channels.^21^ In this case the protein is embedded into a membrane-like lipid bilayer structure which is assembled on top of the scanning surface. Scanning is therefore performed for the combined membrane protein system. We have implemented the *Membrane tool* to roughly mimic such a situation. In a chosen molecular orientation the user can add a solid double layer of adjustable width which is placed parallel to the scanning surface at a specified distance. Those parameters can be chosen to approximately take into account the relative position of the lipid bilayer for the investigated transmembrane protein.

In Fig. 5 we provide a demonstration of the *Membrane Tool* using the PDB structure of the SthK ion channel (PDB 6CJQ). Simulated scanning was performed from the extracellular and the opposing intracellular side. The orientation of corresponding structures was conveniently chosen in the molecular viewer (Fig. 5, middle panel). A proper placement of the double layer with width 4*nm* was obtained using the perspective shown in the *Front View* window (Fig. 5, left panel). Its distance to the scanning surface was adjusted such that the layer surrounds the known transmembrane region of SthK.

**Figure 5:**
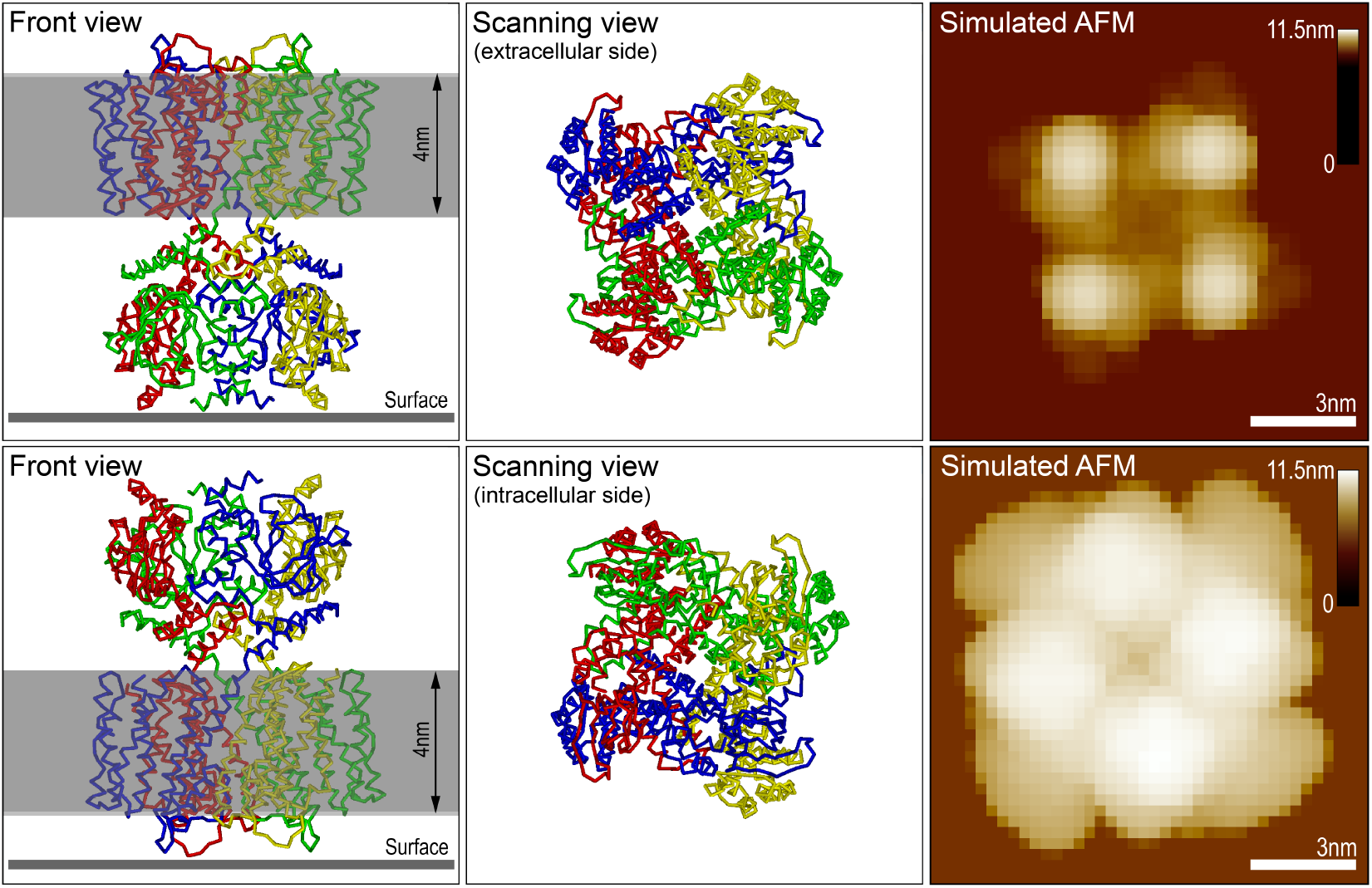
Demonstration of the Membrane Tool. Simulated scanning of the SthK ion channel (PDB 6CJQ) from the extracellular side (top row), and the intracellular side (bottom row). Channel structures in the orientations in which scanning is performed from the top are shown in the middle panel. The corresponding front view perspectives (left panel) display the placement of the solid double layer. On the right side the simulated AFM images are shown together with the respective color bars of detected height for the channel double-layer system.

### Further improvements in BioAFMviewer 2.0

Since the BioAFMviewer software is already used by many AFM groups worldwide, and valuable response helping to improve its functionality has been received, we use this opportunity to mention some of the important improvements we implemented in the BioAFMviewer 2.0 version. The loading of large PDB files, displaying the molecular structures, and, computing the corresponding simulated AFM images, has been highly accelerated to enable a more efficient handling of large protein structures. The integrated molecular viewer now also provides the option to view the loaded structure in the *Backbone View*, in addition to the standard Van-der-Waals (VdW) view. This representation helps in the recognition of secondary structure motifs when biomolecules are compared and fitted to AFM images. However, the user should be aware that simulated scanning is obviously always performed for the atomistic VdW representation. For the graphical representation of the simulated AFM image we included an option to upload a customized color scale according to which topography heights are translated into displayed colors.

A comprehensive explanation of all BioAFMviewer features, including a guide on how to use the fitting and topography tools is provided in an extended manual.

## Discussion

We report the novel BioAFMviewer-Toolbox and provide applications to reconstruct 3D atomistic structures of biomolecules from limited-resolution AFM surface scans. As demonstrated, the implemented topography analysis tools further allow to validate experimental observations. Despite apparent simplifications (see^15^), simulated AFM scanning can remarkably well reproduce experimental images. Its implementation within the BioAFMviewer software, combined with the developed analysis tools, provides the convenient platform for the broad Bio-AFM community to employ the enormous amount of available structural data to facilitate the interpretation and understanding of experimental observations.

We remark that rigid-body fitting of structures to AFM images was also recently presented in,^22^ were a search method similar to our global search was applied. There, it was additionally aimed to infer information on the AFM tip shape from experimental data.

## Material and Methods

### Fitting biomolecular structures to AFM images

#### Quantitative comparison of simulated and experimental AFM images

During the optimization process the agreement of simulated to target AFM image was scored taking into account a variety of quantitative similarity measures.

The image correlation coefficient is based on the pair-wise comparison information encoded in the pixels. It was computed as 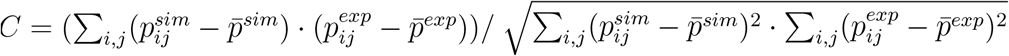, where the summation is performed over all image pixels, 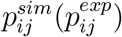 is the information of a pixel at position (*i, j*) in the simulated (experimental) image and, 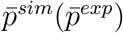 denote average pixel information. The above correlation coefficient provides a standard measure to quantitatively compare two images.

On the other side one can compute a different correlation coefficient which neglects the pixel information relative to the average and instead considers just absolute values, i.e., 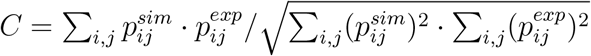.

We also employed the image RMSD and MAE, which quantify the root mean square deviation and mean absolute error of pair-wise pixel information, respectively. They were computed as 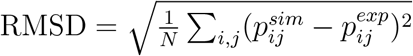, and 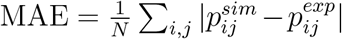, respectively.

We note that pixel information *p*_*ij*_ refers to the color intensity of a pixel, or, alternatively, to real measured height values (if provided for the experimental target image).

#### Details of Global Search

For unbiased sampling of the structure loaded in the molecular viewer, the search space of molecular orientations was obtained by executing rigid-body rotations around all three spatial axes within the coordinate system of our molecular viewer in discrete steps. It was given by the set of discrete orientations 𝒪_*ijk*_ = (*α*_*i*_, *β*_*j*_, *γ*_*k*_) characterized by three angles *α*_*i*_, *β*_*i*_, *γ*_*i*_ = *i* · Ω, where Ω is the angular grid spacing. This spacing parameter can be specified by the user, such that 360° is a multiple of Ω. If, for example, Ω = 10° is chosen, the number of sampled structures in the fitting procedure would be 36^3^ = 46656.

The user can chose to run the global search optimization with either of provided correlation scores, or combinations of them used in parallel. After completed fitting, the user can access the results conveniently from a switch panel, upon which the corresponding fitted molecular structure together with the simulated AFM graphics is displayed in their separate windows, ready to be visually compared to the target experimental AFM image.

#### Details of Local Search

Local search optimization samples structures in an iterative process of small amplitude rigidbody rotations followed by selection, to eventually identify a single molecular structure best fitting the target AFM image. Starting from the initial structure with orientation 𝒪^ini^ = (*α*^ini^, *β*^ini^, *γ*^ini^), a next candidate structure with changed orientation O^cand^ = (*α*^ini^ + *δα, β*^ini^ + *δβ, γ*^ini^ + *δγ*) is obtained by allowing a rigid-body rotation with angle changes *δα, δβ*, and *δγ*. Those angle changes were randomly drawn from a gaussian distribution with center zero and a width value that can be specified by the user. The new candidate structure was then evaluated by scoring the agreement of simulated to target AFM image. Only if the candidate structure improved the agreement it was accepted to replace the initial orientation, i.e. 𝒪^cand^ = 𝒪^ini^, while otherwise it was rejected. The next iteration cycle starts either from the improved fit, or samples other structures around the previous one. The fitting process is terminated when the improvement reached a sufficient level of saturation. It can be noted, that the implementation of the local search procedure is a simplified variant of the Metropolis algorithm of optimization. By selecting only candidate structures which improve agreement scores, the applied method provides an efficient approximation to identify the best fitting structure in the neighborhood of the initial molecular orientation.

#### Application to Hs-AFM images

For each of the 15 selected experimental AFM surface scans of SthK, fitting was based on simulated scanning of the molecular structure (6CJQ) with a fixed tip shape having sphere radius *R* = 2nm and cone-half angle 10°. The chosen spatial resolution corresponded to the experimental value of 0.33nm/pixel. Fitting consisted of applying *Global Search* with a grid spacing of Ω = 40° followed by *Quick Fit* with a variance of 1nm. In Supporting Information T1 we provide a list of similarity scores of simulated to target AFM image obtained for the performed fittings.

For fitting of the actin filament (Figs. 3,4) we used the atomic model Actin model.pdb provided in reference.^24^ The scanning parameters were *R* = 1.5nm, 10° and 0.77nm/pixel. Fitting was obtained by applying the *Quick Fit* function with a variance of 1nm, starting from a *by-hand-chosen* molecular conformation.

### Experimental high-speed AFM images

The SthK channel 2D lattice was obtained from imaging SthK proteins reconstituted in bilayer membranes that were adsorbed on a mica surface. Imaging was performed at a spatial resolution of 0.33nm/pixel. Image tilt correction and horizontal line-by-line leveling as described in^23^ was additionally conducted.

The image of actin was obtained from scanning a phalloidin-stabilized filament on a Mica surface that was treated with (3-aminopropyl)triethoxysilane (APTES). The total scanning area of 60 × 60 nm^2^ was monitored over a grid of 78 × 78 pixels^2^, which defined the spatial resolution we applied for simulated scanning.

## Acknowledgement

We are most grateful to Noriyuki Kodera from the high-speed AFM lab at NanoLSI (Kanazawa University) for his support of the BioAFMviewer software project and his advice in all aspects related to experiments. We also thank him and Noriki Katayama for providing the hs-AFM image of the actin filament.

## Supporting Information Available

**Table T1:** Similarity scores of simulated to target AFM image for fitting of the SthK channel atomic structure to the 15 selected hs-AFM surface scans.

| **SthK AFM**<br>**scan** | **DCC** | **ICC** | **RMSD** |
| --- | --- | --- | --- |
| 1 | 0.94 | 0.89 | 17.91 |
| 2 | 0.85 | 0.78 | 13.99 |
| 3 | 0.90 | 0.80 | 19.32 |
| 4 | 0.89 | 0.86 | 18.32 |
| 5 | 0.97 | 0.94 | 15.77 |
| 6 | 0.87 | 0.80 | 18.85 |
| 7 | 0.94 | 0.84 | 33.51 |
| 8 | 0.89 | 0.76 | 23.94 |
| 9 | 0.96 | 0.93 | 18.82 |
| 10 | 0.87 | 0.58 | 26.93 |
| 11 | 0.95 | 0.92 | 21.64 |
| 12 | 0.94 | 0.86 | 15.68 |
| 13 | 0.94 | 0.82 | 24.20 |
| 14 | 0.92 | 0.91 | 15.65 |
| 15 | 0.96 | 0.90 | 15.97 |

